# Winner’s curse correction and variable thresholding improve performance of polygenic risk modeling based on genome-wide association study summary-level data

**DOI:** 10.1101/034082

**Authors:** Jianxin Shi, Ju-Hyun Park, Jubao Duan, Sonja Berndt, Winton Moy, Kai Yu, Lei Song, William Wheeler, Xing Hua, Debra Silverman, Montserrat Garcia-Closas, Chao Agnes Hsiung, Jonine D Figueroa, Victoria K Cortessis, Núria Malats, Margaret R Karagas, Paolo Vineis, I-Shou Chang, Dongxin Lin, Baosen Zhou, Adeline Seow, Keitaro Matsuo, Yun-Chul Hong, Neil E. Caporaso, Brian Wolpin, Eric Jacobs, Gloria Petersen, Alison P. Klein, Donghui Li, Harvey Risch, Alan R. Sanders, Li Hsu, Robert E. Schoen, Hermann Brenner, MGS (Molecular Genetics of Schizophrenia) GWAS Consortium, GECCO (The Genetics and Epidemiology of Colorectal Cancer Consortium), The GAME-ON/TRICL (Transdisciplinary Research in Cancer of the Lung) GWAS Consortium, PRACTICAL (PRostate cancer AssoCiation group To Investigate Cancer Associated aLterations) Consortium, PanScan and PanC4 Consortium, The GAMEON/ELLIPSE Consortium, Rachael Stolzenberg-Solomon, Pablo Gejman, Qing Lan, Nathaniel Rothman, Laufey T. Amundadottir, Maria Teresa Landi, Douglas F. Levinson, Stephen J. Chanock, Nilanjan Chatterjee

**Author notes:** Correspondences to: Jianxin Shi and Nilanjan Chatterjee.

## Abstract

Recent heritability analyses have indicated that genome-wide association studies (GWAS) have the potential to improve genetic risk prediction for complex diseases based on polygenic risk score (PRS), a simple modelling technique that can be implemented using summary-level data from the discovery samples. We herein propose modifications to improve the performance of PRS. We introduce threshold-dependent winner’s-curse adjustments for marginal association coefficients that are used to weight the SNPs in PRS. Further, as a way to incorporate external functional/annotation knowledge that could identify subsets of SNPs highly enriched for associations, we propose variable thresholds for SNPs selection. We applied our methods to GWAS summary-level data of 14 complex diseases. Across all diseases, a simple winner’s curse correction uniformly led to enhancement of performance of the models, whereas incorporation of functional SNPs was beneficial only for selected diseases. Compared to the standard PRS algorithm, the proposed methods in combination led to notable gain in efficiency (25-50% increase in the prediction R^2^) for 5 of 14 diseases. As an example, for GWAS of type 2 diabetes, winner’s curse correction improved prediction R^2^ from 2.29% based on the standard PRS to 3.10% (P=0.0017) and incorporating functional annotation data further improved R^2^ to 3.53% (*P*=2χ10^-5^). Our simulation studies illustrate why differential treatment of certain categories of functional SNPs, even when shown to be highly enriched for GWAS-heritability, does not lead to proportionate improvement in genetic risk-prediction because of non-uniform linkage disequilibrium structure.

## Introduction

Large genome-wide association studies (GWAS) have accelerated the discovery of dozens or even hundreds of common single nucleotide polymorphisms (SNPs) associated with individual complex traits and diseases, such as height^1; 2^, body mass index^3^ and common cancers (e.g., breast^4^ and prostate^5^ cancers). Although individual SNPs typically have small effects, cumulative results have provided insight about underlying biologic pathways and for some common diseases like breast cancer have yielded levels of risk-stratification that could be useful as part of prevention efforts^6^. Analyses of GWAS heritability using algorithms such as GCTA^7; 8^ have shown that common SNPs have the potential to explain substantially larger fraction of the variation of many traits.

The future yield of GWAS studies, for both discovery and prediction, depends heavily on the underlying effect-size distribution (ESD) of susceptibility SNPs^9; 10, 6^. A number of alternative types of analyses of ESD now point towards a polygenic architecture for most complex traits, in which thousands or even tens of thousands of common SNPs, each with small estimated effect sizes together can explain a substantial fraction of heritability^11; 12^. Mathematical analyses of power indicates that because of the polygenic nature of complex traits, future studies will need large sample sizes, often by an order of magnitude higher than even some of the largest studies to date, for improving accuracy of genetic risk-prediction^10; 11^. Nevertheless, for current datasets, there remains an opportunity to develop more efficient algorithms for improving the models.

Available algorithms for polygenic risk score (PRS) prediction models have varying degrees of complexity. The simplest of these methods, widely implemented in large GWAS, selects SNPs based on a threshold for the significance of the marginal association test-statistics and then the cumulative weighting of these SNPs by their estimated marginal strength of association is applied^13^. The threshold for SNP selection can be optimized to improve the predictive performance in an independent validation dataset. For a number of traits with large GWAS sample sizes, it has been shown that an optimally selected threshold can improve risk prediction compared to that based on the genome-wide significance threshold used for discovery^14^. A number of newer methods involving the joint analysis of all SNPs using sophisticated mixed-effect modeling techniques have recently been developed and may lead further increases in model performance^15-17^.

In this report, we propose simple modifications to the widely used PRS modeling techniques using only GWAS summary-level data. Drawing from the lasso^18^ algorithm, we propose a simple threshold dependent winner’s curse adjustment for marginal association coefficients that can be used to weight the SNPs in PRS. Second, to exploit external functional knowledge that might identify subsets of SNPs highly enriched for association signals, we consider using multiple thresholds for SNPs selection based on group membership and identify an optimal set of thresholds through an independent validation dataset. We demonstrated the value of our new method using summary-level results from large GWAS across a spectrum of traits, some with available independent validation datasets to assess the performance of these methods. Available resources, such as annotation databases, expression and methylation quantitative trait locus (QTL) analyses were employed to identify groups of SNPs that are likely to be enriched with the trait of interest. We evaluated the utility of this information for risk-prediction for respective outcomes. We also report on the performance of new algorithm using simulation studies that incorporate realistic genetic architecture, linkage disequilibrium pattern and enrichment factor for underlying functional SNPs.

## Material and Methods

### PRS construction

Let *Z_m_*, *P_m_*, 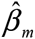, and 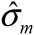 (*m* = 1, …, *M*) denote the univariate *Z*-statistics, the two-sided *P-*values, the estimated association coefficients and their standard deviations available as part of summary-level results for *M*SNPs from a GWAS. We assume that each genotypic value is normalized to have mean zero and unit variance and that 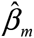 is rescaled to correspond to the normalized genotypic values. Let *g_im_* be the genotype of SNP *m* for subject *i*. The simplest and most popular form of the PRS for GWAS has the form

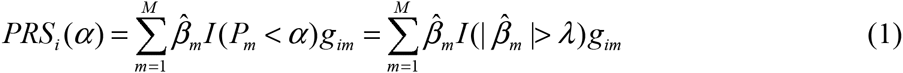

where the threshold *α* for the *P*-values, or equivalently *λ* = Φ^−1^(1 – α/2)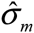 in the β-scale (Appendix A), can be chosen to optimize the predictive performance of PRS in an independent validation dataset. Here, *I*(·) is an indicator function and Φ(·) is the cumulative density function of the standard normal distribution.

Motivated from the simplification of the popular machine learning algorithm lasso^18^ in the orthonormal case, we propose considering a lasso-type thresholding for constructing PRS in the form

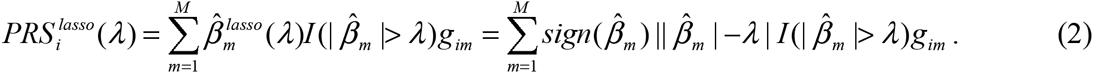

The adjustment of the association coefficient by the threshold parameter in the form of a location-shift can be viewed as a “winner’s curse” bias correction due to nature of the selection of the SNPs. We also considered a more formal approach to winner’s curse bias-correction^19^ by maximizing a conditional likelihood 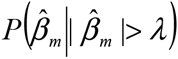 (Appendix A). Let 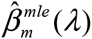 be the maximum likelihood estimate of *β_m_* based on the conditional likelihood. We propose a PRS as

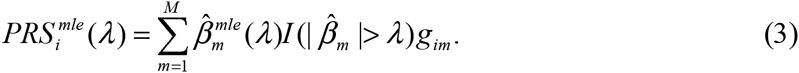

It is, however, critical that for selection of the optimal threshold parameter, bias correction is performed simultaneously with SNP selection for different values of the threshold parameters. We have previously studied the theoretical power for use of such lasso-type winner’s curse correction for developing PRS when SNPs are independent and concluded that under realistic polygenic architecture this simple correction has the potential to improve predictive performance of PRS^11^. The performance of such an algorithm in real GWAS data, where independent SNPs need to be selected after linkage disequilibrium (LD)-filtering, has not been evaluated.

Information from various functional studies, annotation databases and GWAS from various traits is increasingly available to allow identification of subset of SNPs that can be considered to have potential high-prior probability for association with a given trait. Various types of enrichment analyses, whether based on distribution of summary-level statistics^20^ or on more advanced heritability-partitioning analyses^21; 22^, have shown empirical evidence of strong enrichment of GWAS association signals for different categories of SNPs which represent only a relatively small fractions of all GWAS SNPs. However, very few systematic studies have examined whether and how such enrichment information can be utilized to improve models for genetic risk prediction. We consider a simple modification to PRS to explore this issue. We assume that the set of *M*SNPs can be partitioned into two mutually exclusive groups, *S_1_* and *S_2_*, where *S_1_* represents a relatively small subset representing “high-prior” SNPs (referred to as HP) and the second group *S_2_* represents the remainder of the GWAS SNPs (referred to as “low-prior” SNPs or LP) that can be considered part of an “agnostic” search. We allow differential treatment of the SNPs in the PRS:

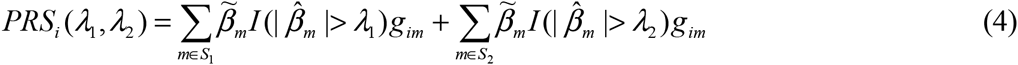

and select the optimal set of threshold parameters based on independent validation dataset(s). We refer to the PRS selecting SNPs with two separate thresholds as two-dimensional PRS or 2D PRS. Here, 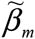 can be chosen as the original estimate 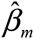, the lasso-type correction 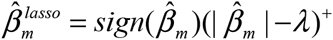 or the MLE correction, 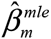. The PRS in (1), (2) and (3) using a single threshold is referred to as 1D PRS.

Following analytic techniques similar to those derived for 1D PRS^11^, we can characterize the theoretical predictive performance of 2D PRS and the corresponding optimal set of thresholds based on the genetic architecture parameters of the two sets of SNPs assuming independence (Appendix B). Using simulation studies, we study performance of the method with realistic LD pattern among SNPs.

### LD-pruning and LD-clumping

The performance of PRS is typically improved if genetic markers are pruned for LD^23^. LD-pruning procedures that ignore GWAS P-values frequently prune out the most significant SNPs and may reduce performance. Instead, we use the LD-clumping procedure implemented in PLINK^23^ that chooses the most significant SNP from a set of SNPs in LD guided by GWAS *P*-values. After LD-clumping, no SNPs with physical distance less than 500kb have LD r^2^ ≥ 0.1.

### Expanding HP SNP set through LD

Suppose *S_1_* is a given HP set defined based on external annotation data (see section *Annotation datasets)*. Any SNP in high LD with a SNP in *S_1_* is also considered to be an HP SNP. Thus, we expanded *S_1_* by including all SNPs that were in high LD (r^2^ ≥ 0.8) with any SNP in the original *S_1_*.

### Simulation Scheme

We performed simulations to evaluate the performance of six PRS prediction methods: 1D and 2D PRS without winner’s curse correction and with lasso/MLE winner’s curse correction. To make simulations realistic in terms of the distribution of minor allele frequencies (MAF) and LD, we simulated quantitative traits with specific genetic architecture by conditioning on the genotypes of a lung cancer GWAS^24^, which had 11,924 samples of European ancestry and 485,315 autosomal SNPs after quality control. The simulation scheme is summarized in the following steps:

(1) We performed LD-pruning implemented in PLINK so that no SNPs within 500kb were in LD at threshold r^2^ = 0.1. After LD-pruning, *M* = 53,163 autosomal SNPs (denoted as *S*) were left.
(2) Denote *S*_1_ as the putative HP SNP set and *S*_2_ = *S*\*S*_1_ as the LP SNP set. We selected a set of 5000 “causal” SNPs (denoted as *C*) from the pruned SNP set *S*. If *C* is randomly selected, *i.e., S*_1_ is not enriched with causal SNPs, we expect | S_1_ ∩*C* |=| *C* ∥ *S*_1_ | /*M*SNPs overlapping between *S*_1_ and *C*. Thus, we defined the enrichment fold change for *S*_1_ as

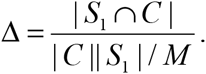 The enrichment fold change Δ ranged from 2 to 4 in simulations.
(3) We simulated quantitative traits according to 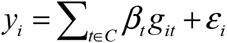, where *β_t_s* were simulated independently from a Gaussian mixture distribution β_t_ ∼ *πN*(0, 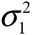) + (1 – *π*)*N*(0, 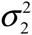) with *π* = 0.1. Here, 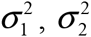 and *Var(ε_j_)* were scaled so that *Var(y_i_)* = 1. The phenotypic variances explained by the two components were 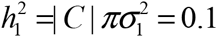 and 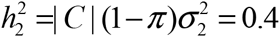. We assume the same effect-size distribution for both HP and LP causal SNPs, but the proportions of causal SNPs are higher in the former than the later group. Under this assumption, Δ also reflects the ratio of heritability explained at a per SNP basis in the HP set compared to LP set.
(4) We randomly selected 10,000 samples as a discovery set and 1,924 as a validation set. We performed GWAS association analysis for all 485,315 autosomal SNPs in the discovery sample. The summary statistics were used to calculate PRS for each sample in the validation sample. The prediction *R*^2^ was calculated as *max_λ_ cor*^2^(*PRS_i_(λ), y_i_*) for 1D PRS methods and max_λ_1_, λ_2__ *cor*^2^(*PRS_i_(λ_1_, λ_2_), y_1_*) for 2D PRS methods. We repeated the simulation 50 times for each set of parameters and report the average prediction *R*^2^. Recently, Finucane et al.^22^ reported the heritability explained by common SNPs in multiple functional categories for 17 traits. Interestingly, they found that common SNPs located in regions that are conserved in mammals^25^ accounted for about 2.6% of total common SNPs but explained approximately 35% of total heritability in average across these traits, suggesting a 13.5-fold enrichment. Thus, we were motivated to investigate whether SNPs related with the conserved regions (CR) may be useful for 2D PRS methods. We downloaded the CR annotations (see Web Resources), identified common SNPs located in any CR and also identified their LD SNPs with *r*^2^ ≥ 0.8. These SNPs are referred as CR-SNPs, which were used as HP *S*_1_ in simulations. We found 9,940 CR-SNPs overlapping with the 53,163 LD-pruned SNPs. To investigate whether specific genomic locations of CR-SNPs influence the performance of 2D-PRS, we also performed simulations using a set *S*_1_ of random SNPs that has the same size and associated heritability as the CR-SNPs.

### GWAS datasets for risk prediction

The information for GWAS data sets and functional annotation data are summarized in Tables S1A and S1B.

## WTCCC GWAS data

The Wellcome Trust Case Control Consortium^26^ (WTCCC) data consisted of two control data sets (1958 Cohort samples and NBS control samples) and seven diseases: bipolar disorder (BD), coronary artery disease (CAD), Crohn’s disease (CD), hypertension (HT), rheumatoid arthritis (RA), Type 1 diabetes (T1D) and Type 2 diabetes (T2D). Since we analyzed T2D using a much larger discovery sample, we did not analyze the T2D data in WTCCC. Because cases and controls were genotyped in different batches, differential errors between cases and controls might cause a serious overestimate of the risk prediction. Thus, we performed very rigorous quality control (QC) by removing duplicate samples, first or second degree relatives, samples with missing rate greater than 5% and non-European samples identified from EigenStrat^27^ analysis. For each disease, we excluded SNPs with MAF<5%, missing rate >2%, missing rate difference >1% between cases and controls or *P*_HWE_<10^-4^ in the control samples. After QC, we had 2,928 controls, 1,817 BD cases, 1,878 CAD cases, 1,729 CD cases, 1,934 HT cases, 1,894 RA cases and 1,939 T1D cases. For each PRS method and each disease, we estimated the prediction R^2^ by five-fold cross-validation.

### Three cancer GWAS with individual genotype data

We analyzed three cancer GWAS with individual level genotype data: the bladder cancer^28; 29^ GWAS of European ancestry including 5,937 cases and 10,862 controls, the pancreatic cancer GWAS^30^ of European ancestry (after excluding samples with Asian or African ancestry) including 5,066 cases and 8,807 controls, and the Asian non-smoking female lung cancer GWAS^31^ with 5,510 cases and 4,544 controls. After QC, the bladder cancer GWAS had 463,559 autosomal SNPs and the Asian lung cancer GWAS had 329,703 autosomal SNPs. The pancreatic cancer GWAS included samples from three studies that used different genotyping platforms. For convenience, we analyzed 267,935 autosomal SNPs that overlapped in all three platforms. The prediction performance was evaluated using ten-fold cross-validation.

### Five large GWAS with summary statistics and independent validation samples

For T2D, we downloaded the summary statistics of the DIAGRAM (DIAbetes Genetics Replication And Meta-analysis) consortium^32^ with 12,171 cases and 56,862 controls for 2.5 million SNPs imputed to the Hapmap2 reference panel. We also downloaded the GERA (Genetic Epidemiology Research on Adult Health and Aging) GWAS data of European ancestry with 7,131 T2D patients and 49,747 samples without T2D (but may have other medical conditions, e.g., 27.4% with cancers, 25.4% with asthma, 25.4% with allergic rhinitis and 12.4% with depression). Although these non-T2D samples were not perfect healthy controls, we found that most of the genome-wide significant SNPs in DIAGRAM could be replicated in GERA (data not shown). We randomly selected 5,631 T2D patients and 48,247 non-T2D subjects from GERA as discovery set, performed association analysis adjusting for top 10 PCA scores and meta-analyzed with the summary statistics from DIAGRAM for 353,196 autosomal SNPs overlapping between the two studies. The resulting summary statistics were used to build PRS risk models, which were validated in the remaining 1500 T2D patients and 1500 non-T2D subjects in GERA.

The PGC2 (Psychiatric Genetics Consortium) schizophrenia GWAS meta-analysis consisted of 34,241 cases and 45,604 controls^33^. Summary statistics were obtained by meta-analyzing all PGC2 schizophrenia GWAS except the MGS^34^ (Molecular Genetics of Schizophrenia) subjects of European ancestry. The summary statistics were used to build PRS models, which were validated in MGS samples with 2,681 cases and 2653 controls.

The TRICL (Transdisciplinary Research in Cancer of the Lung) GWAS consortium consisted of 12,537 lung cancer cases and 17,285 controls^35; 36^. We performed meta-analysis using TRICL samples excluding the samples from the PLCO^24^ (Prostate, Lung, Colon, and Ovary Cohort Study) study. The summary statistics based on 11,300 cases and 15,952 controls were used to build risk models, which were validated in the PLCO lung GWAS samples with 1,237 cases and 1,333 controls.

For colorectal cancer, we performed meta-analysis for the GECCO (Genetics and Epidemiology of Colorectal Cancer Consortium)^37^ GWAS data after excluding the PLCO GWAS data. The PLCO samples were genotyped using two different genotyping platforms with different marker densities: one had approximately 500K SNPs and the other had only 250K SNPs. Thus, we first imputed the genotypes to the Hapmap2 reference panel using IMPUTE2^38^ and selected SNPs with imputation *r^2^* ≥ 0.9 for risk prediction. The discovery sample consisted of 9,719 cases and 10,937 controls from 19 studies. The PLCO validation sample had 1,000 cases and 2,302 controls.

The summary statistics for prostate cancer were obtained from the PRACTICAL (PRostate cancer AssoCiation group To Investigate Cancer Associated aLterations) consortium and The GAME-ON/ELLIPSE (Elucidating Loci Involved in Prostate Cancer Susceptibility) Consortium with samples from populations of European, African, Japanese and Latino ancestry^5^. The discovery samples consisted of 38,703 cases and 40,796 controls after excluding the NCI Pegsus GWAS samples with 4,600 cases and 2,941 controls, which were used for validation. We analyzed 536,057 autosomal SNPs after QC that overlapped between the validation and the discovery sample summary statistics.

#### Annotation datasets

For many traits, GWAS risk SNPs have been reported to show enrichment for eQTLs, methylation QTLs (meQTLs) and cis-regulatory elements (CREs). In addition, recent studies have reported extensive genetic pleiotropy across diseases and traits, e.g. psychiatric diseases^39; 40^, schizophrenia and cardiovascular-disease risk factors, including blood pressure, triglycerides, low- and high-density lipoprotein, body mass index (BMI) and waist-to-hip ratio (WHR)^41^. Thus, we defined the HP SNP set *S*_1_ using eQTL SNPs (referred to as eSNPs) in blood, tissue specific eSNPs and meQTL SNPs (referred to as meSNPs), SNPs related with CREs (referred to as CRE-SNPs), SNPs related with genomic regions conserved across mammals (referred to as CR-SNPs) and SNPs identified by pleiotropic analyses (referred to as PT-SNPs). We expanded each SNP set by including LD SNPs with *r*^2^ ≥ 0.8 in the local 1M region for each SNP. Here, LD was calculated based on the genotype data of relevant ancestry in The 1000 Genomes Project^42^.

<u>*eSNPs and meSNPs*</u>: Blood *cis*-eSNPs were identified from two large-scale eQTL studies in European populations. One study involved a transcriptome sequencing project of 922 subjects^43^ and the other involved a microarray study of 5,311 subjects^44^. Because of its very large sample size, the second study had the power to detect eSNPs with even tiny effect sizes which may not have meaningful functional importance. Thus, we included eSNPs with association *P*-value <10^-6^ with any gene in the *cis* region in the second study. For both Asian and European lung cancer GWAS data, we used eSNPs^45^ and meSNPs^46^ based on lung tissues. For T2D, we used eSNPs^47^ and meSNPs^48^ based on adipose tissues. Furthermore, detected *trans*-SNPs are much fewer than *cis*-SNPs and the replication rate of *trans*-eSNPs was much lower than *cis*-SNPs^47^, suggesting that including *trans*-SNPs would be unlikely to improve risk prediction. Thus, we did not include *trans*-SNPs.

<u>*CRE-SNPs:*</u> CREs are regions of noncoding DNA regulating the transcription of nearby genes. SNPs located in CREs may change the binding of specific transcription factors and thus the expression of the target genes. Typically, CREs are identified through ChIP-Seq experiments of histone modifications. We downloaded “peak” data (each peak represents one CRE) of specific sets of histone methylation markings, acetylation markings and DNase I hypersensitive sites (DHSs) from the ROADMAP project website for relevant cell lines. For each identified CRE (‘peak’), we identified common SNPs with MAF>1%. For prostate cancer, we used the ChIP-Seq data for H3K27Ac and the transcription factor TCF7L2^49^ to define HP SNP sets.

<u>*PT-SNPs:*</u> The summary statistics for height^1; 2^, BMI and obesity^3; 50^, WHR^51^, waist circumference (WC)^51^, hip circumference (HIP)^51^ were downloaded from the GIANT consortium website. The summary statistics for GWAS meta-analysis of cardiovascular-disease risk factors^52^, including triglycerides (TG), low-density lipoprotein (LDL) and high-density lipoprotein (HDL), were also used for 2D PRS.

We investigated whether or not each tentative HP SNP set was enriched for GWAS associations by examining the quantile-quantile (QQ) plot, which was made for HP SNPs vs. LP SNPs after LD-clumping. The SNP sets not enriched for GWAS associations were not expected to improve risk prediction in 2D PRS. Thus, for each disease, we only included HP SNP sets for 2D PRS when they showed strong enrichment in QQ plots. Interestingly, blood eSNPs were enriched for almost all diseases. CR-SNPs showed modest enrichment for majority of the diseases. Thus, blood eSNPs and CR-SNPs were used for 2D PRS for all diseases. In addition, eSNPs and meSNPs derived in lung tissues were enriched in lung cancer GWAS of both European and Asian ancestry. The SNPs related in enhancer and active promoter regions (characterized by H3K4me3, H3K9-14Ac, H3K36me3, H3K4me1, H3K9ac and H3K9me3) were enriched for GWAS associations but SNPs related with the repressive regions (characterized by H3K27me3) were not. Thus, we included SNPs related with these enhancer and active promoter regions for 2D PRS. DHS SNPs were not strongly enriched and thus were excluded. Recently, we have shown significantly shared genetic component between lung cancer and bladder cancer risk^53^. Thus, we also used HP SNPs derived based on lung tissues or cell lines for predicting bladder cancer risk. Furthermore, we found that SNPs identified through pleiotropic analysis were enriched in multiple diseases. For example, SNPs with *P*-value <0.001 in GWAS of height, HDL, LDL, TC, TG, WC, obesity, HIP and T2D were enriched in lung cancer GWAS. Because our 2D PRS methods required a relatively large number of HP SNPs to achieve improvement, we combined the SNPs with *P*-value <10^-3^ (or 10^-2^) in at least one trait into a HP SNP set referred as PT-0.001 (or PT-0.01).

#### Testing the statistical significance of improvement for risk prediction

For WTCCC and three cancer GWAS data sets with individual genotype data, we used K-fold cross-validation to estimate prediction R^2^. Here, *K*=5 for WTCCC data and *K*=10 for cancer GWAS data. We were interested in testing whether the prediction of a new PRS method was significantly better than that of the standard 1D PRS defined in equation (1). For the *i^th^* crossvalidation, we denote 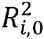 as the maximum prediction for the standard 1D PRS optimized across *P*-value thresholds, 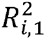 as the maximum prediction for a new PRS method optimized across all *P*-value thresholds for 1D PRS and all pairs of *P*-value thresholds for 2D PRS. We defined 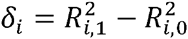 and estimated its variance as 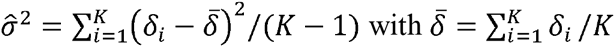. We calculated the statistic 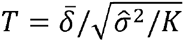 and evaluated its significance using the *t*-distribution. For the five diseases with independent validation samples, we used bootstrap to estimate the variance of the R^2^ estimates to test significance^42^.

## Results

### Theoretic investigation of 2D PRS performance assuming independent SNPs

Figure 1A shows the theoretically-derived AUC for a binary trait based on 1D PRS and 2D PRS without applying a winner’s curse correction. For all PRS models, the AUC increases with the sample size of the discovery dataset. The 2D PRS can improve the 1D PRS in which the magnitude depends on the sample size in the discovery sample and also the enrichment of the HP SNPs. Figure 1B shows the optimal *P*-value thresholds for including SNPs that maximize the prediction of 2D PRS. The optimal P-value threshold for including HP SNPs is more liberal than that for LP SNPs and the difference diminishes as the training sample size becomes very large.

**Figure 1.**
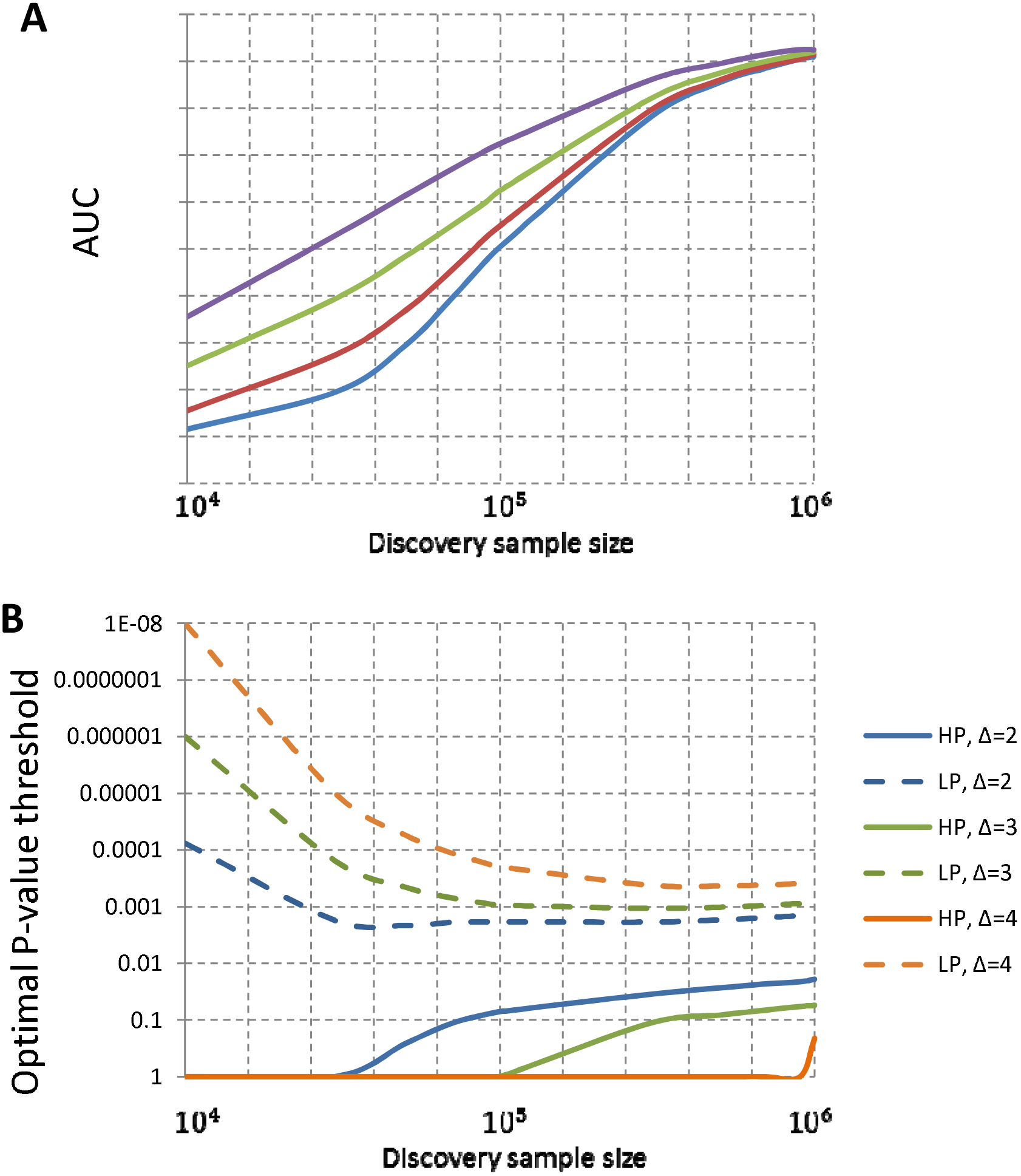
Theoretic investigation of prediction performance and optimal thresholds for SNP selection in 2D PRS. The theoretic calculation assumes *M* = 53,163 independent SNP, of which 5,000 are causal for a binary trait, similar to simulation studies. The high-prior (HP) SNP set has 5,000 SNPs and the low-prior (LP) SNP set has 48,163 SNPs. Δ is the enrichment fold of HP SNPs in the causal SNP set. (A) The prediction AUC for 1D PRS and 2D PRS. (B) The optimal *P*-value thresholds for including HP and LP SNPs in 2D PRS. For both plots, x-coordinate is the discovery sample size, assuming equal number of cases and controls.

### Polygenic risk prediction of T2D

Figure 2A presents the 1D PRS results for T2D. The standard 1D PRS without winner’s curse correction had a prediction R^2^=2.29% by including SNPs with P≤2×10^-3^. The winner’s curse correction improved R^2^ to 3.10% using the lasso-type correction and 2.67% using the MLE correction.

**Figure 2.**
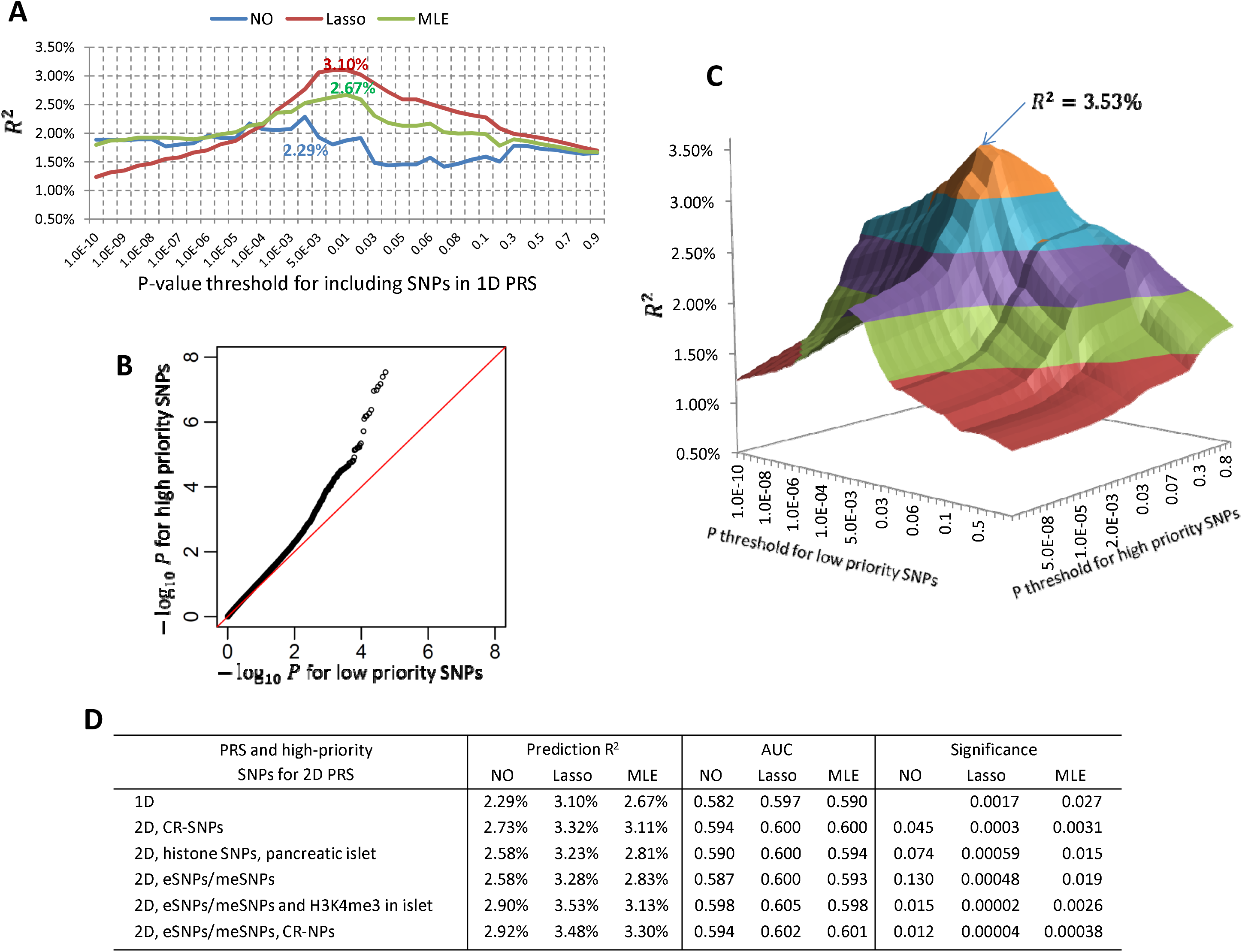
Genetic risk prediction for type-2 diabetes. PRS models were built based on the summary statistics from a meta-analysis of DIAGRAM consortium and GERA data (17,802 cases and 105,109 controls in total) and validated in independent 1500 cases and 1500 controls in GERA. (A) Prediction R^2^ (observational scale) for 1D PRS with or without winner’s curse correction. “NO”: no winner’s correction for association coefficients; “Lasso”: regression coefficients were modified by a lasso-type correction; “MLE”: association coefficients were modified by maximizing a likelihood function conditioning on selection. (B) Quantile-quantile plot for –log_10_(*P*) for high priority (HP) SNPs v.s. low priority (LP) SNPs. SNPs were pruned to have pairwise *r*^2^ ≤ 0.1. Here, the HP SNPs were eSNPs/meSNPs in adipose tissue or SNPs related with the H3K4me3 mark in pancreatic islet cell line with data downloaded from the ROADMAP project. The HP SNPs were strongly enriched in the discovery data. (C) Prediction R^2^ for 2D PRS with lasso-type winner’s curse correction. The SNP set was the same to (B). The best prediction (R^2^ =3.53%) was achieved when we included HP SNPs using criterion *P* ≤ 0.03 and HP SNPs with *P* ≤ 0.005. (D) The prediction R^2^, the area under the curve (AUC) and the significances for testing whether an alternative PRS was better than the standard 1D. The Nagelkerke R^2^ values were reported in Tables S4.

Next, we investigated whether functional annotation could further improve risk prediction. We considered CR-SNPs, eSNPs and meSNPs in adipose tissue, and SNPs related with different histone marks and their combinations as HP SNP sets. These SNPs were enriched in T2D GWAS, exemplified by the QQ plot in Figure 2B for a HP SNP set comprising of eSNPs/meSNPs in adipose tissue and SNPs related with H3K4me3 in the pancreatic islet cell line. Note that the SNPs have been pruned to have LD r^2^≤0.1, so the observed enrichment was unlikely due to an artifact related to extensive LD. Figure 2C illustrates how the prediction R^2^ of a 2D PRS depends on the *P*-value thresholds for the HP and LP SNPs. The prediction R^2^ was maximized using a more liberal *P*-value threshold 0.03 for HP SNPs and a more rigorous threshold 0.005 for LP SNPs. This optimal 2D PRS had 8,018 HP SNPs and 2,033 LP SNPs.

Figure 2D reports the prediction R^2^, AUC and the significance for testing of whether an alternative PRS method could improve the standard 1D PRS. The best predictions were achieved by the 2D PRS with lasso-type correction: R^2^=3.48% using eSNPs/meSNPs and CR-SNPs and R^2^=3.53% using eSNPs/meSNPs and H3K4me3 SNPs in pancreatic islet cell line (52.0% and 54.1% efficiency gain compared to 2.29% using standard 1D PRS, respectively). These improvements were statistically significant compared to the 1D standard PRS (*P*=0.00002 and 0. 00004, respectively). Of note, the recently developed method LD-pred^54^ that models the LD information only slightly improved prediction R^2^ from 2.47% to 2.73% (10. 5% efficiency gain) using DIAGRAM summary statistics as discovery.

### Results for WTCCC data

The prediction R^2^ values for six diseases in WTCCC data are reported in Figure 3A. The AUCs and Nagelkerke R^2^ are summarized in Table S2. Optimal thresholds for SNP selection are in Table S3. The lasso-type winner’s curse correction improved the 1D PRS predictions for CD (6.65% to 8.22%), RA (7.24% to 8.60%) and T1D (18.2% to 18.5%) and was slightly better than the MLE winner’s curse correction. The 2D PRS improved the prediction for CD (6.65% to 7.71% using blood eSNPs). Combining functional data and lasso-type correction gave a prediction R^2^=8.75% for CD (31.6% efficiency gain over the standard 1D PRS). Note that our method of winner’s curse correction together 2D PRS performed at least as well as the standard 1D PRS. However, because of the small sample size in the validation sample, the improvements were not statistically significant.

**Figure 3.**
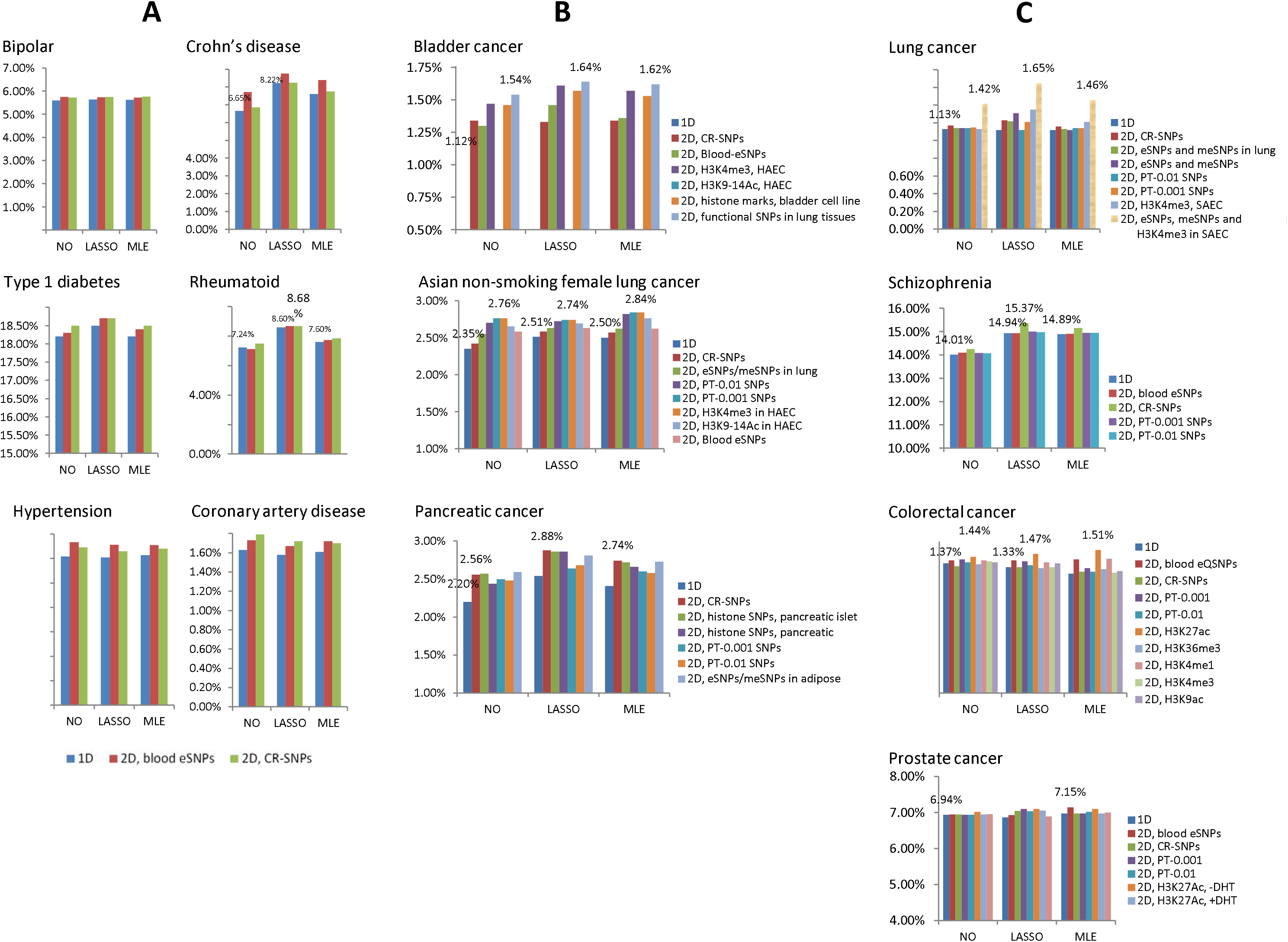
Comparison of polygenic risk prediction methods for 13 complex diseases. For all figures, the y-coordinate is the prediction R^2^ in the observational scale. “1D” denotes 1D PRS; “2D, blood eSNPs” denotes 2D PRS using blood eSNPs as high-prior SNP set. In the x-axis, “NO” denotes PRS without winner’s curse correction; “LASSO” and “MLE” denote lasso-type and MLE winner’s curse correction, respectively. (A) Prediction R^2^ values for six diseases in WTCCC data, estimated based on five-fold cross-validation. (B) Prediction R^2^ values for three GWAS of cancers, estimated based on ten-fold cross-validation. (C) Prediction R^2^ values for four complex diseases estimated based on independent validation samples.

### Results for three cancer GWAS with individual genotype data

Results are summarized in Figure 3B (prediction R^2^), Table S4 (AUC and Nagelkerke R^2^), Table S5 (*P*-value for testing significance of improvement) and Table S6 (optimal thresholds for SNP selection). The standard 1D PRS achieved an R^2^ =1.12% for bladder cancer, 2.35% for Asian nonsmoking female lung cancer and 2.2% for pancreatic cancer, indicating the difficulty of genetic risk prediction for these cancers. The lasso-type correction improved the 1D PRS for all three cancers: R^2^ from 1.12% to 1.29% for bladder cancer, 2.35% to 2.51% for Asian female nonsmoking lung cancer and 2.20% to 2.54% for pancreatic cancer. Our 2D PRS methods further improved the prediction although the various annotation datasets gave different improvement. For bladder cancer, the greatest efficiency gain (R^2^ =1.64%, 46.4% efficiency gain over the standard 1D PRS and 27.1% efficiency gain over the 1D PRS with lasso-type correction) was achieved with the SNPs related to the lung tissue/cell line expression data (eSNPs, meSNPs, H3K4me3 SNPs in SAEC), which performed slightly better than the SNPs related with histone marks in bladder cell line (R^2^ =1.46%). For non-smoking female Asian lung cancer, the 2D PRS incorporated with PT-0.001 SNPs or H3K4me3 SNPs in HAEC improved R^2^ to 2.84%. For pancreatic cancer, the 2D PRS incorporated with CR-SNPs, SNPs related with histone marks of pancreatic islet and adipose eSNPs/meSNPs improved prediction R^2^ by approximately ∼30% compared with the standard 1D PRS. Many of the improvements over the standard 1D PRS were statistically significant (Table S5), e.g., P=0.025 for 2D PRS with H3K4me3 SNPs in HAEC for bladder cancer, P=0.025 for 2D PRS with PT-0.001 SNPs for Asian lung cancer and P=0.047 (0.023, 0.023) for 2D PRS with CR-SNPs (PT-0.001, PT-0.01 SNPs) for pancreatic cancer.

### Results for four large-scale summary-statistics datasets

Prediction results are reported in Figure 3C (prediction R^2^), Table S7 (AUC and Nagelkerke R^2^), Table S8 (*P*-values for testing whether improvements were significant), Table S9 (optimal p-value thresholds for SNP selection in 2D PRS) and Figure S2. For lung cancer, the standard 1D PRS had an R^2^=1.13%. The best prediction R^2^=1.65% (a 46.0% efficiency gain compared with the standard 1D PRS) was achieved by lasso-corrected 2D PRS with eSNPs/meSNPs in lung tissues, blood eSNPs and SNPs related with H3K4me3 in SAEC. To achieve this prediction accuracy, the optimal *P*-value threshold for the 2D PRS should be 0.008 for HP SNPs and 5×10^-6^ for LP SNPs. However, the improvement was not statistically significant. For schizophrenia, the lasso-type correction improved 1D PRS R^2^ from 14.01% to 14.94%; the 2D PRS with CR-SNPs further improved the R^2^ to 15.37% and the improvement was highly statistically significant (P=3.2×10^-10^). The optimal p-value threshold was 0.6 for CR-SNPs and 0.1 for other SNPs in 2D PRS with lasso-type correction. For CRC and prostate cancer, neither winner’s curse correction nor 2D PRS improved prediction.

### Simulation results

The simulation results are summarized in Figure 4. First, the winner’s curse corrections, both lasso-type correction and MLE correction, slightly improved prediction in most if not all simulations and in particular improved more for the 1D PRS than the 2D PRS. We also observed that the two winner’s curse correction methods performed similarly. Second, if HP SNPs were chosen randomly in the LD-pruned SNP set and were strongly enriched for causal SNPs, the 2D PRS methods substantially improved the prediction over the 1D PRS methods. As expected, the improvement increased quickly with the enrichment fold change Δ. Without winner’s curse correction, 1D PRS had R^2^=1.38% and 2D PRS improved R^2^ to 2.13% for Δ=2, 2.86% for Δ=3 and 4.22% for Δ=4. Consistent with theoretical analysis assuming independent SNPs (Figure 1B), the optimal *P*-value threshold for HP SNPs was more liberal than that for LP SNPs (Table S10).

**Figure 4.**
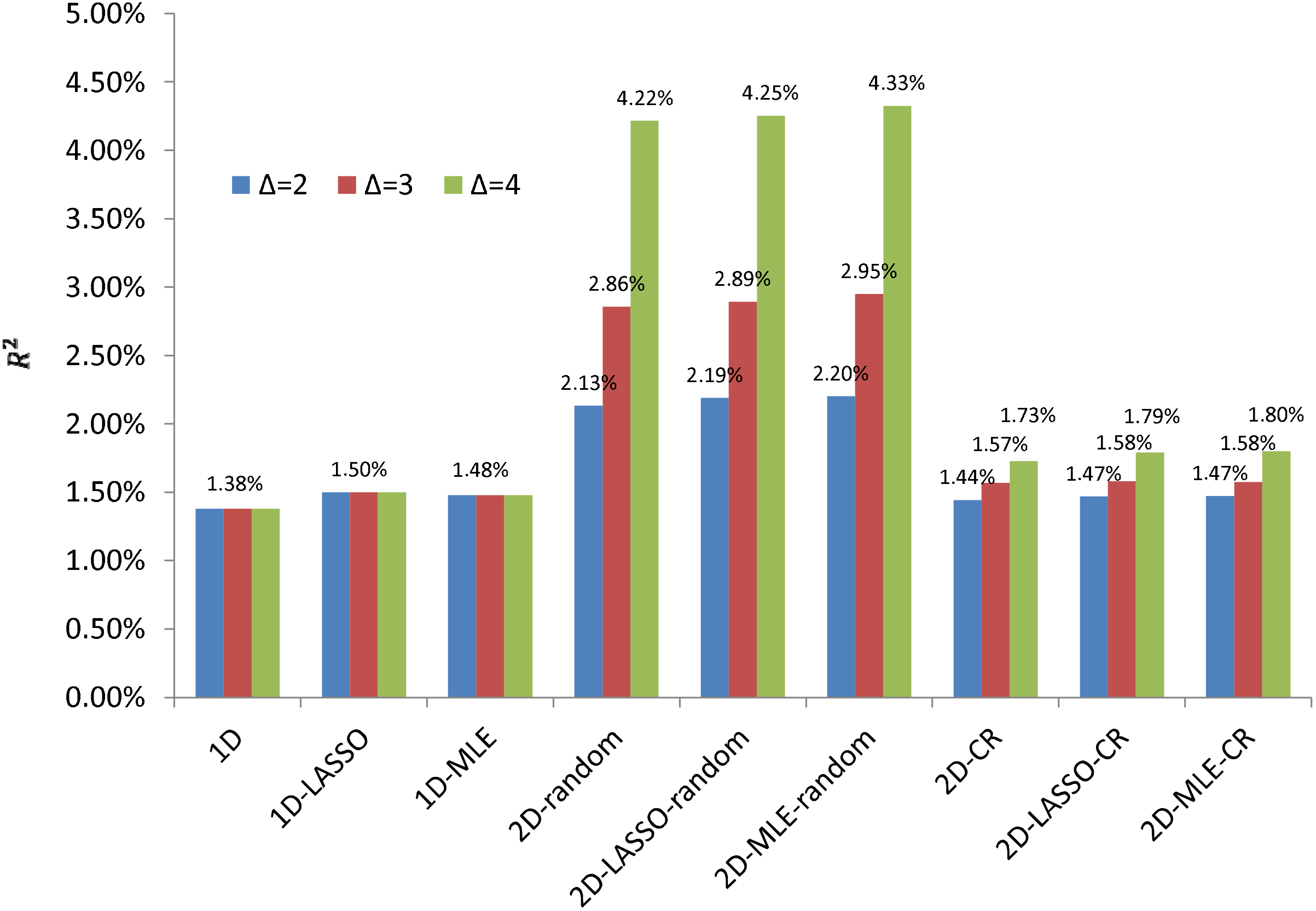
Simulation results for comparing polygenic risk prediction methods and different high priority SNP sets. Quantitative traits were simulated conditioning on the genotypes of LD-pruned SNPs in lung cancer GWAS with 10,000 discovery samples and 1,924 validation samples. For each simulation, we used 5,000 causal SNPs and 9,940 high priority (HP) SNPs (either randomly selected or the SNPs related with conserved regions). Δ denotes the enrichment fold change of the HP SNP. In the x-axis, “1D” denotes 1D PRS without winner’s curse correction; “1D-LASSO(MLE)” denotes 1D PRS with lasso-type (MLE) correction; “2D-random” indicates 2D PRS with HP SNP sets randomly selected from the LD-pruned SNPs in the genome; “2D-CR” indicates 2D PRS using SNPs in conserved regions as HP SNPs.

However, when we used CR-SNPs as the HP SNPs, the improvement of 2D PRS was less compared to the simulations with randomly selected HP SNPs, even with the same enrichment fold change. As a numerical example, when Δ=4, the 2D PRS method without winner’s curse correction improved R^2^ from 1.38% to 1.73% for CR-SNPs as HP SNPs while from 1.38% to 4.22% for random HP SNPs. To investigate whether the difference was caused by different local LD structure, for each SNP, we counted the number of SNPs located less than 1Mb from the given SNP and had *r^2^* ≥ 0.8 with the SNP in the 1000 Genomes Project. For 9,940 CR-SNPs used for our simulations, the average number of LD SNPs is 22.4 (median=12) while the average number is 6.4 (median=2) for non-CR SNPs. See also the histograms in Figure S1. Thus, CR-SNPs are enriched in regions with strong LD and may suggest a possible explanation why CR-SNPs (and other functional categories with similar LD structure) may not lead to improvement in risk prediction as much as would be expected based on enriched heritability.

## Discussion

Our study demonstrates that the predictive performance of GWAS PRS models can be improved based on a combination of a simple adjustment to the threshold levels of SNP selection and weights of selected SNPs. The degree of gain, however, is not uniform and depends on multiple factors, including the genetic architecture of the trait, sample size of the discovery sample set, degree of enrichment of association in selected set of “high-prior” SNPs and the linkage disequilibrium patterns of these SNPs with the rest of the genome.

The simple winner’s curse correction of SNP weights using the lasso-type method leads to an improvement in performance of PRS uniformly across all studied diseases. For some diseases, such as type-2 diabetes (Figure 2 and Table S7) or Crohn’s disease (Figure 3 and Table S2), this correction alone led to notable improvement in the performance of PRS. The optimal weighting of SNPs would depend on the true effect size distribution of the underlying susceptibility SNPs. Lasso-type weights can be expected to be optimal under a double exponential distribution^18; 55^, and it is possible that the weighting could be improved further under alternative models of effect-size distribution. It is, however, encouraging that irrespective of what might be the true effect-size distribution, which is likely to vary across the diseases of study, our simple lasso-correction improves over the standard PRS methodology without adding any additional computational complexity.

The effect of using various threshold levels for different functional categories of SNPs on the performance of the model varied by disease as well as the functional annotation of external data sets employed in our analytical approach. After adjustment with lasso-type weights, the use of two-dimensional threshold based on prioritized SNPs led to notably higher values of R^2^ for lung cancer in Caucasians (increase by 46% using eSNPs, meSNPs and SNPs related with H3K4me3 in SAEC as high priority set), bladder cancer (increased by 27.1% using high priority SNPs in lung tissue or cell lines), type-2 diabetes (increased by 13.9% using eSNPs, meSNPs and SNPs related with H3K4me3 mark in islet cell line) and pancreatic cancer (increased by 10.6% using SNPs related with histone modification marks in pancreatic or islet cell lines). Consistent with theoretical expectations, for each of the traits, the optimal thresholds selected were more liberal for the associated category of high-prior SNPs than those for complementary set.

Our simulation study illustrated how the improvement in performance of the PRS model due to differential treatment of certain categories of SNPs is modest even when these SNPs have been categorized to be highly enriched for heritability^21^. For example, recent heritability partitioning analysis has identified SNPs in conserved DNA regions, representing 2.6% of the genome, to be highly enriched for GWAS heritability for many diseases (explaining 35% heritability on average). Our theoretical calculations suggest that if only independent SNPs are analyzed, use of a subset of SNPs similarly enriched for heritability is expected to yield much higher improvement in the performance of the model (Figure 1). Our simulation studies showed that a similarly large gain is expected even in the presence of naturally occurring LD pattern if these SNPs are selected randomly from the genome. However, when we simulated high-prior SNPs based on the exact location of the CR-SNPs, the improvement was modest, within the range of observed data. The CR-SNPs represent a highly unusual linkage disequilibrium pattern in that they are in high degree of LD with an unusually large number of neighboring SNPs (Figure S1).

In the future, more detailed and accurate assessment of the functional annotation of SNPs should improve performance of PRS models. Our method requires only simple modifications to the standard PRS algorithm and can thereby be used to rapidly evaluate the effectiveness of many alternative strategies. In the current study, we used physical location information pertaining to histone marks to define high-priority SNP. However, a SNP located in histone marks does not necessarily cause the variation in histone binding. Thus, a more reasonable approach is to identify genetic variants associated with histone variation across subjects in order to define high-priority SNP sets. These types of histone QTLs have recently been reported in small-scale studies based on HapMap samples^56; 57^. We expect that histone QTL SNPs identified in future large-scale tissue specific studies might be more informative for risk prediction.

We have investigated the performance of the various algorithms using criteria that reflect how much of the variability of the observed outcomes can be explained by the PRS in the validation dataset. For clinical applications of risk-models, however, it is important to evaluate whether models are well calibrated that is to what extent they can produce unbiased estimates of risk for individuals with different SNP profiles. Earlier studies have noted that the standard PRS can be mis-calibrated and additional calibration steps may be needed when applying PRS in a clinical setting. In this regard, we find that a winner’s curse correction can alleviate calibration bias of the standard PRS, but substantial residual bias remains in some situations (Table S11). The regression relationship between overall PRS and disease status can be estimated based on a relatively small validation sample and can also be used to re-scale PRS for producing calibrated risk estimates.

We used several different metrics for evaluating the potential impact of an improved PRS for risk-stratification. The percentage gain in prediction R^2^ due to improved PRS is substantial for several diseases. For these diseases, the impact of an improved PRS on overall discriminatory performance of the models is noticeable but small (increase in AUC value between 1-2%). However, even a modest increment in AUC value can lead to identification of substantially higher fraction of individuals who are at the tails of risk distribution and hence likely to consider clinical decisions (Table S12).

A limitation of our method is that we use stringent LD-pruning for creating sets of independent SNPs. However, this may result in loss of predictive power of models as SNPs in moderate or low LD may still harbor independent association signals. The LD-pred^54^ method has been proposed to better account for correlated SNPs in building PRS using GWAS summary-level data and has been shown to lead to improved performance over standard PRS for some diseases such as schizophrenia. The LD-pred method also uses a specific form of prior distribution for obtaining “shrunken” estimates of the regression coefficients for the SNPs in the model. Although we did not make direct comparisons, it appears that the LD-pred method gains over standard PRS by improving the accounting for correlation between risk SNPs. In contrast, in our algorithm, which used stringent LD pruning, the gain in performance over the standard PRS mainly came from the lasso-type winner’s curse correction and the use of variable thresholds to account for HP and LP SNPs. Thus it is possible that in the future the complementary strengths of the algorithms can be combined to develop more powerful PRS.

In conclusion, we have proposed a set of simple methods for constructing PRS for genetic risk prediction using GWAS summary-level data. The proposed methods are computationally not onerous and yet show a noteworthy gain in performance. A major strength of our study is that we evaluated the proposed methods across a large number of scenarios reflecting a spectrum of underlying genetic architectures for different complex diseases, sample size of the study and available functional annotation. These studies and additional simulations provide comprehensive insights to promises and limitations of genetic risk prediction models in the near future.

## Appendices

### Appendix A: Winner’s curse correction

#### Lasso-type winner’s curse bias correction estimator

Suppose that for a given SNP, we have the two-sided P-value P_i_, the regression coefficient 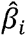 its standard deviation 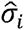 and the *Z*-statistic 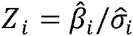. The SNP is included in a risk prediction model if P_*i*_ ≤ *α* or equivalently |Z_i_| ≥ Φ^-1^(1 – α/2) or equivalently 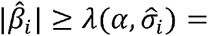 Φ *–11–α/2σi*. Here, Φ() is the cumulative distribution function of *N(0,1)*. The lasso-type shrinkage estimator conditioning on *P_i_* ≤ *α* is given as 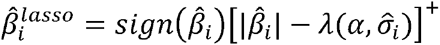. Note that the bias correction depends on the p-value threshold *a* for including SNPs.

#### Maximum likelihood estimator to reduce winner’s curse bias

Following Zhong and Prentice^19^, we assume 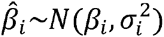 with 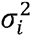 approximated by 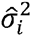. By conditioning on *P_i_* ≤ *α* or equivalently 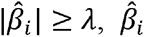 follows a truncated normal distribution with an explicit density function

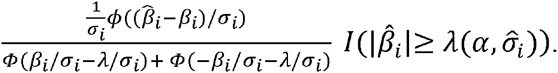

We derived the estimator 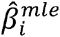 by maximizing the conditional likelihood numerically using R. Again, 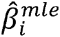 depends on the p-value threshold *α* for including SNPs. For computational efficiency, we pre-calculated 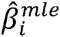 at a required precision for all predefined p-value thresholds.

### Appendix B: Theoretical prediction performance assuming independent SNPs

Suppose that for a given trait of interest *Y*, there are two predefined SNP sets: the high priority (HP) SNP set *S*_1_ and the low priority (LP) SNP set *S*_2_. SNPs have been pruned and are in linkage equilibrium. We assume that *S*_1_ has *M*_1_ independent susceptibility SNPs and *M*_3_ null SNPs while *S*_2_ has *M*_2_ susceptibility SNPs and *M*_4_ independent null SNPs. Following Chatterjee et al.^11^, we assume that the true relationship between outcome *Y* and independent susceptibility SNPs is modeled as follows:

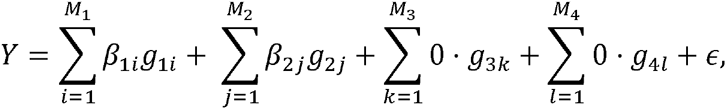

where all *Y* and the genotypic values *g*’s are standardized so that *E(Y)* = 0, *Var(Y)* = 1, *E(g)* = 0 and *Var(g)* = 1, and the error term ∊∼*N*(0, *σ*^2^) and is independent of the genotypic values.

From a discovery GWAS data set of size *N*, we have regression coefficient 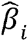 and two-sided p-value *P_i_* for each SNP. We build an additive prediction model by including SNPs in *S*_1_ with *P*-value ≤ *α*_1_ and SNPs in *S*_2_ with *P*-value ≤ *α*_2_:

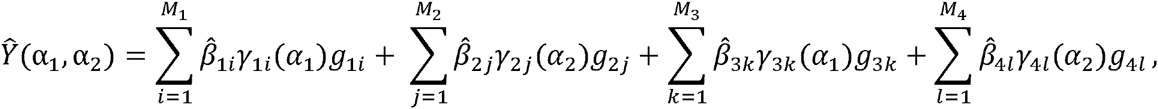

where *γ(α)* = *I(P ≤ α)* with *I*(·) being an indicator function.

The predictive correlation coefficient (PCC) for the predictive model can be expressed as

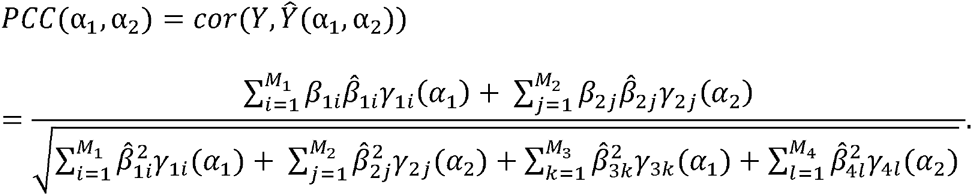

Following Chatterjee et al. (2014), one can verify that PCC follows a normal distribution by the central limit theorem and the strong law of large numbers. Therefore, the expected value of PCC can be approximated as

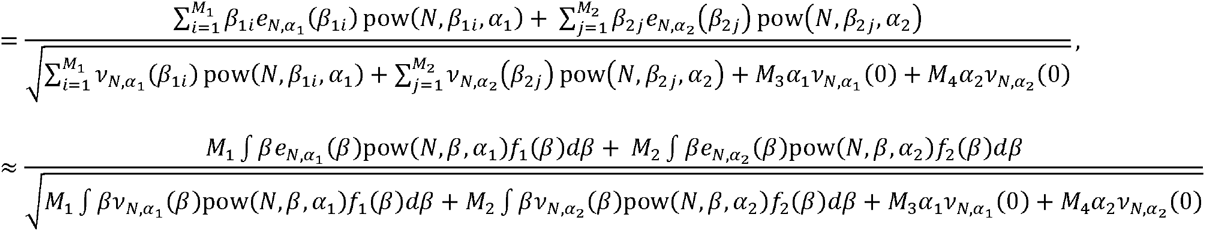

where *e_N, α_(β)* = *E*(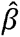|γ(α) = 1), *v_N, α_(β)* = *E*(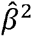|γ(α) = 1), pow(*N, β, α*) is power to detect a SNP with effect size *β* at a significance level α in a GWAS with size *N*, and *f*_1_(·) and *f*_2_(·) are effect-size distributions for HP and LP susceptibility SNPs, respectively.

In our numerical calculations, we assumed that the effect sizes of the susceptibility SNPs in the HP and LP sets followed the same distribution *β∼πN*(0, 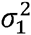) + (1 – *π*)*N*(0, 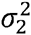), consistent with simulations. We performed grid search to identify the p-value thresholds (α_1;_ α_2_) that maximizes *E*(*PCC*(α_1_, α_2_)). For binary disease outcomes, the area under the curve (AUC) can be expressed as a function of PCC, as shown in Chatterjee et al. (2014).

## Acknowledgements

This study utilized the high-performance computational capabilities of the Biowulf Linux cluster at the National Institutes of Health, Bethesda, MD. (http://biowulf.nih.gov). This study made use of data generated by the Wellcome Trust Case Control Consortium (WTCCC). A full list of the investigators who contributed to the generation of the data is available at www.wtccc.org.uk. Funding for the WTCCC project was provided by the Wellcome Trust under award 076113. J.S. and N.C. were supported by the NIH intramural research program. The TRICL Consortium was supported by NIH grant U19 CA148127. We thank Hilary Kiyo Finucane and Alkes Price for providing the annotation data for conserved DNA regions. We would like to acknowledge all the investigators, their support staff, and their funding support who contributed to GWAS of lung cancer among non-smoking females in Asia, as part of the Female Lung Cancer Consortium in Asia (FLCCA), described in reference 31. We would like to acknowledge all the investigators, their support staff, and their funding support who contributed to GWAS of bladder cancer, described in reference 28 and in reference 29.

## Web Resources

The URLs for data provide herein are as follows:

Annotation for conserved genomic regions: http://compbio.mit.edu/human-constraint/data/gff/

DIAGRAM type 2 diabetes summary statistics, http://diagram-consortium.org/downloads.html

GERA GWAS data; http://www.ncbi.nlm.nih.gov/projects/gap/cgibin/study.cgi?study_id=phs000674.v1.p1

IMPUTE2, https://mathgen.stats.ox.ac.uk/impute/impute_v2.html

Psychiatric Genomic Consortium (PGC2), schizophrenia summary statistics, http://www.med.unc.edu/pgc/downloads

Histone mark and DHS peak data, http://www.roadmapproject.org/

Conserved genomic regions, http://compbio.mit.edu/human-constraint/data/gff/

Height, BMI, WC, WHP, obesity summary statistics from GIANT consortium, http://www.broadinstitute.org/collaboration/giant/index.php/GIANT_consortium

LDL, HDL, TC and triglycerides summary statistics, http://www.broadinstitute.org/mpg/pubs/lipids2010/

eQTL and meQTL in adipos, http://www.muther.ac.uk/Data.html

Blood eQTL, http://genenetwork.nl/bloodeqtlbrowser/

SNAP, http://www.broadinstitute.org/mpg/snap/

Transdisciplinary Research In Cancer of the Lung (TRICL), http://u19tricl.org/

The code for PRS data analysis is available at http://dceg.cancer.gov/tools/analysis/functionalPRS

